# HIPPOCAMPAL THETA OSCILLATIONS DURING CONTROLLED SPEED RUNS ON A TREADMILL

**DOI:** 10.1101/2025.06.17.660140

**Authors:** Alan Michel Bezerra Furtunato, Daniel Marques da Silva, Adriano Bretanha Lopes Tort, Bruno Lobão-Soares, Hindiael Belchior

## Abstract

Hippocampal theta (6–12 Hz) oscillations coordinate neural activity during spatial navigation and are strongly related to locomotion speed. However, recent research has yielded conflicting evidence on whether theta rhythms are primarily modulated by acceleration or instantaneous speed. Moreover, the role of movement transitions—at locomotion onset and offset—has often been overlooked, despite potentially involving distinct dynamics not explained by speed or acceleration alone. Previous studies have rarely controlled for locomotion timing and speed, limiting our ability to dissociate the contributions of speed, acceleration, and movement transitions. To address this, we used a computer-controlled treadmill to induce rat locomotion under three distinct conditions: (a) movement transitions, (b) steady running at constant speed, and (c) locomotion with continuous acceleration. This setup allowed precise spectral analysis of hippocampal theta oscillations across conditions. We found that treadmill-triggered movement transitions produced sustained increases in theta power and transient increases in theta frequency. Upon treadmill stop, theta power decreased slowly, whereas theta frequency dropped rapidly. Steady running elevated both theta power and frequency relative to rest. During constant-speed trials, both metrics increased with speed and remained stable over time. Notably, the acceleration rate itself had no effect on theta power or frequency. Instead, during accelerating trials, theta frequency increased progressively with instantaneous speed, underscoring speed as the primary modulator. In summary, our results show that movement transitions induce distinct, sustained changes in theta power and transient changes in theta frequency, while instantaneous speed—not acceleration—governs hippocampal theta frequency.

**Significance statement:** The precise contributions of movement transitions, speed, and acceleration to hippocampal theta oscillations remain unclear due to confounding factors in freely moving paradigms. To resolve this, we employed a computer-controlled treadmill to systematically isolate each locomotor variable under tightly controlled conditions. Our results demonstrate that movement transitions induce distinct changes in theta power and frequency, and that instantaneous speed—not acceleration—robustly modulates theta frequency across hippocampal subregions. These findings clarify an ongoing debate and refine our understanding of how specific locomotor dynamics shape hippocampal activity during navigation.

## Introduction

Theta (6-12 Hz) oscillations are large-amplitude rhythmic neural activities produced by the hippocampal local field potentials (LFPs). Hippocampal theta oscillations are believed to coordinate network activity across the trisynaptic circuit (CA1, CA3, and the dentate gyrus—DG) and other cortical structures (Colgin, 2013). These oscillations have been extensively linked to spatial navigation, learning, and memory, as well as movement-related processes in rodents and humans (Jarrard, 1993; Oddie and Bland, 1998; Eichenbaum, 2000; Buzsáki, 2005; Buzsáki and Moser, 2013; Rolls et al., 2018; Nuñez and Buño, 2021).

Early research by Vanderwolf and Heron (1964) and Vanderwolf (1969; 1971) reported that hippocampal theta oscillations emerge during movement preparation and voluntary locomotion, while McFarland et al. (1975) further demonstrated that the intensity of voluntary movements evokes proportional changes in theta activity. Over the past decades, studies have consistently shown that locomotion modulates hippocampal theta oscillations, with faster movement often associated with stronger theta power and higher frequency (Bland and Oddie, 2001; Buzsáki, 2005; McNaughton et al., 2006). However, most studies have been conducted in freely moving rodents navigating open fields or linear and circular tracks while executing behavioral tasks, which could introduce major sensorimotor, cognitive, and emotional confounds. Despite these potential confounds, most studies in freely moving rodents have generally reported a positive correlation between locomotion and theta oscillations (Sławińska and Kasicki, 1998; Maurer et al., 2005; Montgomery et al., 2009; Hinman et al., 2011; Bender et al., 2015).

Beyond freely moving conditions, a few studies have imposed different types of locomotion to evaluate their effects on hippocampal theta. For instance, passive locomotion experiments, utilizing apparatuses such as cars and trains, have shown that translational displacement without active body movements also stimulates hippocampal theta, thereby emphasizing the role of sensory inputs in generating this rhythm (Terrazas et al., 2005; Xie et al., 2012; Li et al., 2014; Cei et al., 2014; Drieu et al., 2018). Comparisons between translational and stationary locomotion (e.g., on a wheel) revealed higher theta power and slower theta frequency during stationary conditions, possibly due to lower visuospatial and vestibular stimuli when running in place at equivalent speeds (Safaryan and Mehta, 2021; De Lima and Belchior, 2023). Similarly, head-fixed rats in virtual reality setups exhibited slower theta frequency compared to freely moving conditions, likely due to reduced vestibular stimuli (Safaryan and Mehta, 2021). Comparisons between treadmill and running wheel locomotion showed higher changes in theta frequency at the treadmill and a stronger association between changes in theta frequency and heart rate, suggesting that treadmill locomotion exerts greater emotional arousal (Li et al., 2014).

To further investigate the contributions of locomotion parameters, other studies have attempted to actively control the speed of locomotion to identify specific changes in theta oscillations in response to acceleration and instantaneous running speed. Furtunato et al. (2020) have shown that theta power decreases across consecutive treadmill running trials at the same speed. Kropff et al. (2021) designed an experiment with rats running inside a bottomless car to evaluate the effects of locomotion at constant speeds and during acceleration and deceleration on hippocampal theta oscillations. They concluded that acceleration had a significant influence on theta frequency, with minor effects on theta power. However, Kennedy et al. (2022) later reanalyzed the dataset and found that running speed explained most of the variability in theta power and frequency, suggesting that the original findings might have been influenced by biased speed sampling.

In the present study, we use a speed-controlled treadmill with 6 different running protocols and spectral analysis to disentangle the effects of movement transitions, locomotion speed, and acceleration on theta oscillations obtained from the DG, CA3, and CA1 areas of the dorsal hippocampus. We found that treadmill-triggered movement transitions caused sustained changes in theta power and transient changes in theta frequency, and that, regardless of acceleration, hippocampal theta was reliably modulated by instantaneous locomotion speed.

## Materials and Methods

### Animals

We used five adult male Wistar rats (3 months old; 350-450 g) provided by the Central Animal Facility of the Biosciences Center at the Federal University of Rio Grande do Norte. After electrode implantation, the rats were housed individually in transparent plexiglass cages (19 x 12 x 7 cm) and maintained on a 12h/12h light-dark schedule (lights on at 6 am) with food and water ad libitum. Experiments were conducted between 4 pm and 8 pm. All procedures were approved by the Ethics Committee on the Use of Animals (CEUA/UFRN, permit n° 52/2016) and followed the US National Institutes of Health Guide for the Care and Use of Laboratory Animals (Institute of Laboratory Animal Research, 2011).

### Electrodes

We built microelectrode arrays of 16 electrodes arranged in two 1x8 bundles, with an inter-electrode depth spacing of 200 μm, designed to bilaterally target the dorsal hippocampus (-3.6 mm AP, ± 3.0 mm ML, according to Paxinos and Watson, 2009). We recorded electrophysiological signals from CA1, CA3, and DG areas. Detailed descriptions of microelectrode array manufacture and surgical implantation procedures can be found elsewhere (Neves et al., 2022).

### Experimental protocol

We used an electrical treadmill (AVS Projetos) with belt dimensions of 40 x 13.5 cm. A metallic grid positioned at the back of the belt delivered low-intensity electrical shocks (0.1 to 0.7 mA at 60 Hz for up to 1 second) when animals moved out of the belt. Electrical shocks were applied only during the habituation and training phases, and were not used during the recording sessions. Treadmill activity was controlled by a microcontroller (Arduino Uno) and stored as TTL inputs by the electrophysiological acquisition system.

Animals were initially habituated to the experimenter and the treadmill apparatus for 3 days. On the first day, rats were placed on the treadmill for 30 minutes with both the electrical shock function and the treadmill belt deactivated. On the second day, the electrical shock function was activated, but the treadmill belt remained off. On the third day, rats walked on the treadmill at a low speed (10 cm/s) for 5 minutes after 30 minutes of habituation with the shock function activated.

The experiment was designed for rats to run on a treadmill with controlled accelerations (A1 = 1 cm/s^2^, A2 = 1.5 cm/s^2^, A3 = 2 cm/s^2^) or constant speeds (S1 = 20 cm/s, S2 = 30 cm/s, S3 = 40 cm/s) in the following sequence: A1, A2, A3, S1, S2, S3. This sequence of 6 trial types was repeated 8 times per session day. Each running trial lasted 20 seconds and was interleaved with a 5-second rest interval. Rat 1 performed one session, while Rats 2, 3, 4, and 5 executed four recording sessions each, totaling 17 sessions. Each trial type (three accelerations and three constant speeds) was therefore recorded 136 times (i.e., 8 repetitions x 17 sessions).

### Data collection and analysis

Continuous intracranial electrophysiological recordings were performed using a headstage preamplifier wire coupled to a multi-channel recording system (RHA2116, Intan Technologies). Raw electrophysiological signals were amplified 200x, bandpass filtered between 0.02 Hz and 20 kHz, and sampled at 30 kHz. Electrophysiological recordings and treadmill TTL inputs were synchronized and stored for posterior analysis. Rat 1 was video recorded by a high-definition digital camera (1080 x 720 pixels at 30 frames/s, Logitech C920) positioned perpendicularly to the treadmill.

Electrophysiological analyses were made in MATLAB using customized scripts, built-in functions, and the EEGLAB Toolbox (Delorme and Makeig, 2004). Raw electrophysiological signals were downsampled to 1 kHz and detrended to obtain LFPs. One electrode per hippocampal area/hemisphere was selected based on its anatomical targeting and theta phase profile, similar to Scheffer-Teixeira et al. (2012). Specifically, we filtered the LFP in the theta (6-12 Hz) band using the “eegfilt” function from the EEGLAB Toolbox. The "hilbert" function from the Signal Processing Toolbox was used to obtain the instantaneous theta phase of each LFP, from which we calculated the theta phase difference between the most superficial electrode and all other electrodes in the same hemisphere. Since theta phase reversal is known to occur between CA1/CA3 areas and the DG, close to the stratum radiatum and stratum lacunosum-moleculare (Brankačk et al., 1993, Buzsáki, 2002; Csicsvari et al., 2003; Scheffer-Texeira et al., 2012; Bragin et al., 1995), signals with the lowest phase difference relative to the theta phases exhibited by the most superficial electrode were selected as CA1 electrodes, signals with the highest phase difference (i.e., phase reversal) were selected as DG electrodes, and signals with the lowest phase difference and anatomically positioned within the fourth quartiles of the bundle were selected as CA3 electrodes. Finally, LFP segments from running trials were visually inspected to exclude epochs containing mechanical or electrical artifacts. The resulting number of valid trials was as follows: A1 = 130, A2 = 127, A3 = 118, S1 = 125, S2 = 122, S3 = 107.

Spectral analyses were performed on LFP data from epochs of treadmill runs and adjacent rest intervals. We used the “spectrogram” function (1-s window, 50% overlap) from the Signal Processing Toolbox to obtain the time-frequency decomposition, as shown in Figure 2. Theta peak frequency was defined as the frequency within the 6-12 Hz band with maximum power in the spectrogram. The average theta power was calculated as the mean power between 6 and 12 Hz.

**Figure 1.**
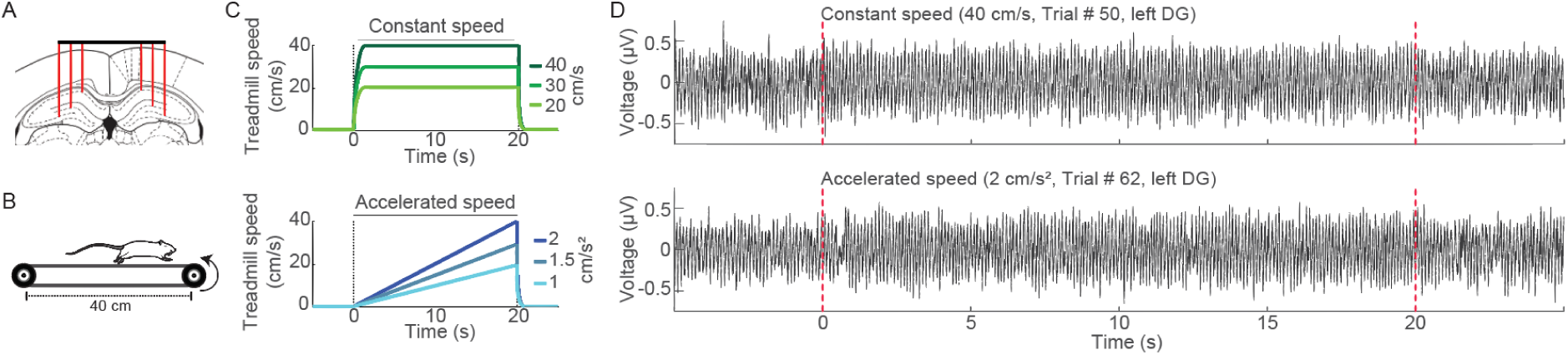
In vivo electrophysiological recordings during treadmill running at constant and accelerating speeds and inter-trial rest intervals. (A) Representative image depicting the anatomical structure and the positioning of electrodes in the DG, CA3, and CA1 areas of the hippocampus. (B) Schematic illustration of the treadmill. (C) Treadmill running protocols at constant speeds of 20, 30, and 40 cm/s (upper) and accelerating speeds of 1, 1.5, and 2 cm/s^2^ (lower). The duration of each running trial was 20 s, with inter-trial rest intervals lasting 5 s. Dashed lines indicate treadmill onset and offset times. (D) Representative LFP recordings from the left DG during a constant speed trial (upper, 40 cm/s) and an accelerating speed trial (lower, 2 cm/s^2^), including the adjacent 5-s rest intervals.

**Figure 2.**
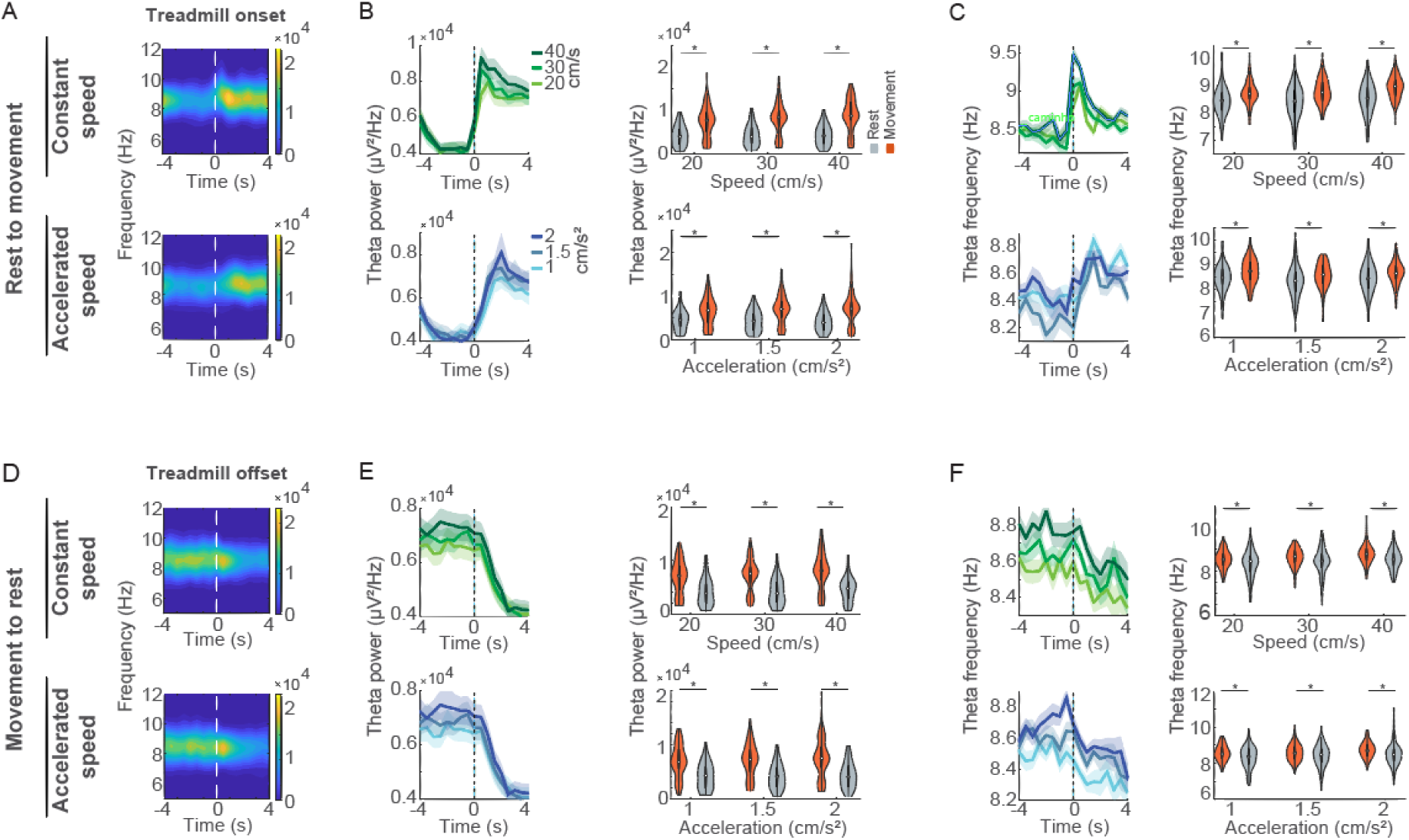
Hippocampal theta power and frequency change dynamically at movement transitions. (A) Average spectrograms of the left DG LFPs aligned to treadmill onset (time 0) during constant speed (20, 30, and 40 cm/s, upper) and accelerating speed (1, 1.5, and 2 cm/s², lower) running trials. (B, left) Theta (6-12 Hz) power around transitions from rest to movement for constant and acceleration trials. Darker green and blue lines indicate higher constant and accelerating speeds, respectively. (B, right) Violin plots show the distribution, median, and interquartile range of theta power for the last 2 s of rest and the first 2 s of running. Asterisks denote p<0.01, Wilcoxon signed-rank sum test. (C) Similar to B, but for theta frequency. (D-F) Similar to A-C, but aligned to treadmill offset (time 0).

We used the “pwelch” function (1-second window, no overlap) from the Signal Processing Toolbox to compute the power spectral density (PSD). To exclude transitions between rest and locomotion, the first 5 seconds of each running trial were discarded from this analysis, resulting in epochs of 15 s. In a separate set of analyses, these 15-s periods were further divided into non-overlapping 5-s blocks. Power spectral estimates were obtained on a trial-by-trial basis for each electrode. The theta power was defined as the average power values in the 6-12 Hz range, and theta peak frequency from its local maximum.

### Statistical analysis

MATLAB (MathWorks) was used for statistical analyses, with significance set at α = 0.05. We used the Shapiro-Wilk test to analyze data normality. Parametric data were analyzed using Student’s t-test or ANOVA, followed by the Tukey-Kramer post-hoc test. Non-parametric data were evaluated through the Wilcoxon signed-rank sum test or the Kruskal-Wallis test, followed by the Tukey-Kramer post-hoc test.

## Results

To evaluate the influence of running speed and acceleration on hippocampal oscillations, we analyzed LFP recordings from the DG, CA3, and CA1 areas of five rats performing a treadmill running task (Figure 1A and 1B). The rats completed consecutive treadmill runs under constant speeds (20, 30, and 40 cm/s) and accelerating speeds (1, 1.5, and 2 cm/s^2^; Figure 1C). Each run lasted 20 s and was followed by a 5-s rest period. Figure 1D displays representative raw DG LFPs during typical rest (<0 s and >20 s) and treadmill running periods (0-20 s) at constant and accelerating speeds.

### Theta oscillations change dynamically at movement transitions

Since previous studies have shown that hippocampal theta oscillations are affected by movement initiation (Whishaw and Vanderwolf, 1973; Vanderwolf and Heron, 1964), we first investigated how movement transitions impact the power and frequency of these oscillations. Average spectrograms aligned to treadmill onset and offset revealed dynamic changes in spectral components (Figure 2A and D). The power and frequency of the theta rhythm increased following the transition from rest to movement in both constant and acceleration trials (Figure 2B and C, left). Notably, the changes in theta power and frequency were larger at the highest treadmill speeds. Consequently, theta changes were stronger (∼20% higher) and more abrupt in constant speed trials compared to accelerating trials. After reaching its maximum, theta power remained elevated at ∼80% of its peak value, while theta frequency returned closer to baseline levels, highlighting different temporal dynamics of these two spectral features during movement initiation. The Wilcoxon signed-rank sum test confirmed that both theta power and theta peak frequency were significantly higher after treadmill onset across all speeds and hippocampal areas (e.g., p < 0.001 for most comparisons, Wilcoxon signed-rank sum test, 2-s epochs before and after treadmill onset; Figure 2B, right, and 2C, right; see Supplementary Table 2-1 for full statistics).

In contrast, transitions from movement to rest induced changes in the opposite direction (Figure 2D). Theta power and frequency returned to baseline after the treadmill offset with similar temporal dynamics for constant and acceleration trials (Figure 2E and F, left). However, decreases in theta frequency exhibited slightly slower temporal dynamics. Theta power and frequency were significantly lower after the treadmill offset for all speed protocols and hippocampal areas (Figure 2E, right, and 2F, right, p < 0.01, Wilcoxon signed-rank sum test; Supplementary Table 2-2).

### Theta power and theta frequency during locomotion vs. rest periods

Next, we compared inter-trial rest intervals and treadmill running periods, excluding movement transitions (i.e., we discarded the first 5 seconds after treadmill onset; Figure 3). Spectral analysis showed an overall increase in broadband power during running (0-250 Hz, Figure 3A, p<0.05, Wilcoxon signed-rank sum test). Theta power significantly increased compared to inter-trial rest intervals across all hippocampal areas (DG, CA3, and CA1) and for both constant and acceleration trials (Figure 3B and Supplementary Table 3-1, p<0.01, Wilcoxon signed-rank sum test). These increases were not due to nonspecific broadband power elevation, as the relative contribution of the theta band to total power also increased significantly (e.g., in the left DG, from 66.46% to 74.68% during constant speed trials, and from 64.63% to 73.21% during acceleration trials, p<0.01, Wilcoxon signed-rank sum test). Similarly, the theta peak frequency was higher during treadmill running than at inter-trial rest intervals (Figure 3C and Supplementary Table 3-1, p<0.05, Wilcoxon signed-rank sum test). However, this effect was stronger during running at constant speeds than at accelerating speeds (Figure 3C, upper and lower, respectively), in which theta peak frequency significantly changed only at the lowest acceleration (i.e., 1 cm/s²; p<0.05, Wilcoxon signed-rank sum test).

**Figure 3.**
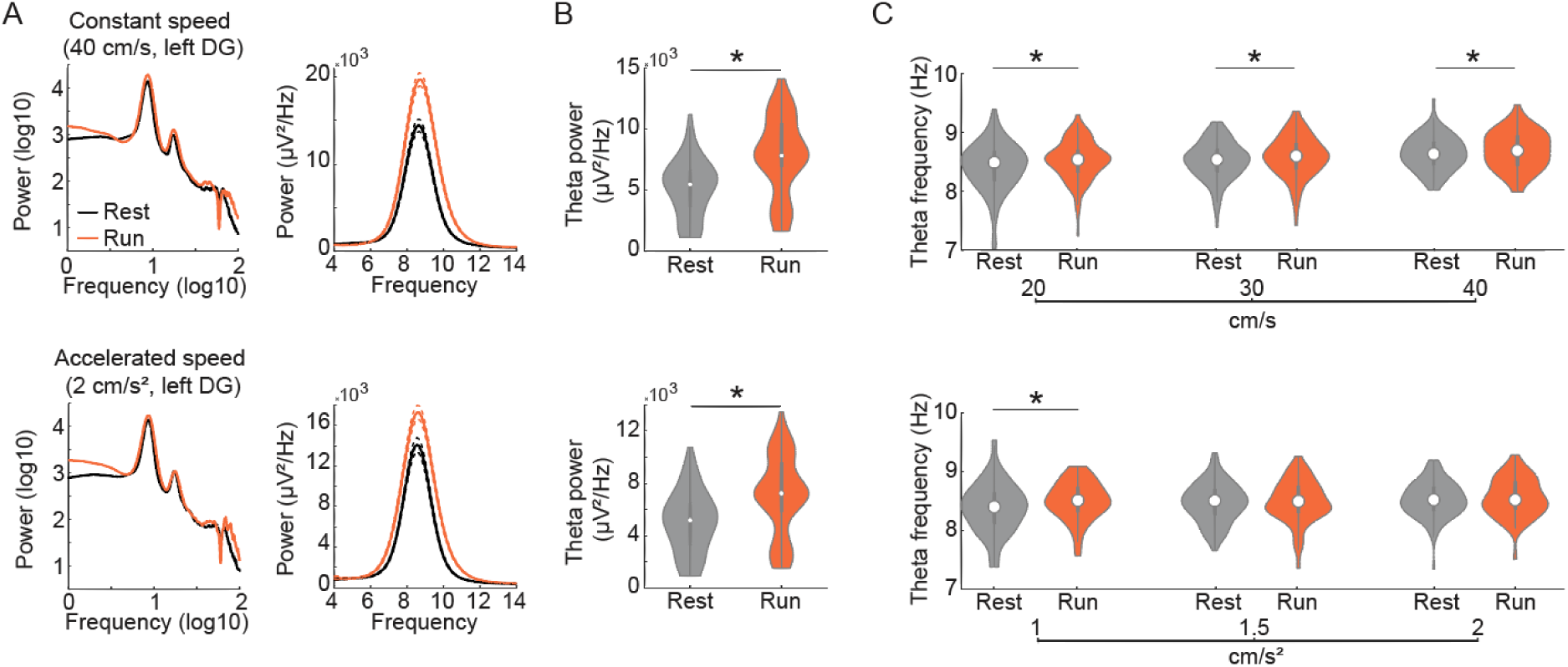
Theta power and peak frequency during treadmill running and at inter-trial rest intervals. (A, left) Broadband LFP power spectra from the left DG during treadmill running (red) and inter-trial rest intervals (black). Spectra are shown for constant (upper, 40 cm/s) and accelerating speed (lower, 2 cm/s²) trials. (A, right) Average power spectra at the theta band; dashed lines represent SEM. (B) Violin plots showing the distribution of theta power during inter-trial rest intervals and treadmill running at constant (40 cm/s, upper) and accelerating speeds (2 cm/s², lower). Asterisk denotes p<0.05, Wilcoxon signed-rank sum test. Similar results were obtained for all treadmill protocols (see Supplementary Table 3-1). (C) Violin plots showing the distribution of theta peak frequency during inter-trial rest intervals and treadmill running at constant (upper) and accelerating speeds (lower). All results in this figure were obtained from LFP segments excluding movement transition periods (see Methods).

### Treadmill running speed affects hippocampal theta power and frequency

Next, we compared theta power and frequency during running at different constant speeds (Figure 4A). To isolate steady-state locomotion, the first 5 seconds following treadmill onset were excluded from the analysis. Figure 4B shows the average power spectra at the theta band, which indicate stronger and faster theta oscillations at higher running speeds. Theta band power significantly increased as running speed increased only in the left DG and right CA3 areas (Figure 4C left, p<0.05 and p<0.01, respectively, Kruskal-Wallis test), while other areas had no significant changes (see Supplementary Table 4-1). In contrast, changes in theta peak frequency were more prominent: theta frequency significantly increased with treadmill running speed across DG, CA3, and CA1 in both hippocampal hemispheres (Figure 4C right, p<0.05, Kruskal-Wallis test). To assess whether theta oscillations varied over the course of a constant-speed run, we divided the 15-s running period at 40 cm/s into three consecutive 5-s blocks (5–10 s, 10–15 s, and 15–20 s; Figure 4D). In this analysis, no significant differences in theta band power or peak frequency were observed across time blocks in any hippocampal area (Figure 4E and F, see also Supplementary Table 4-1).

**Figure 4.**
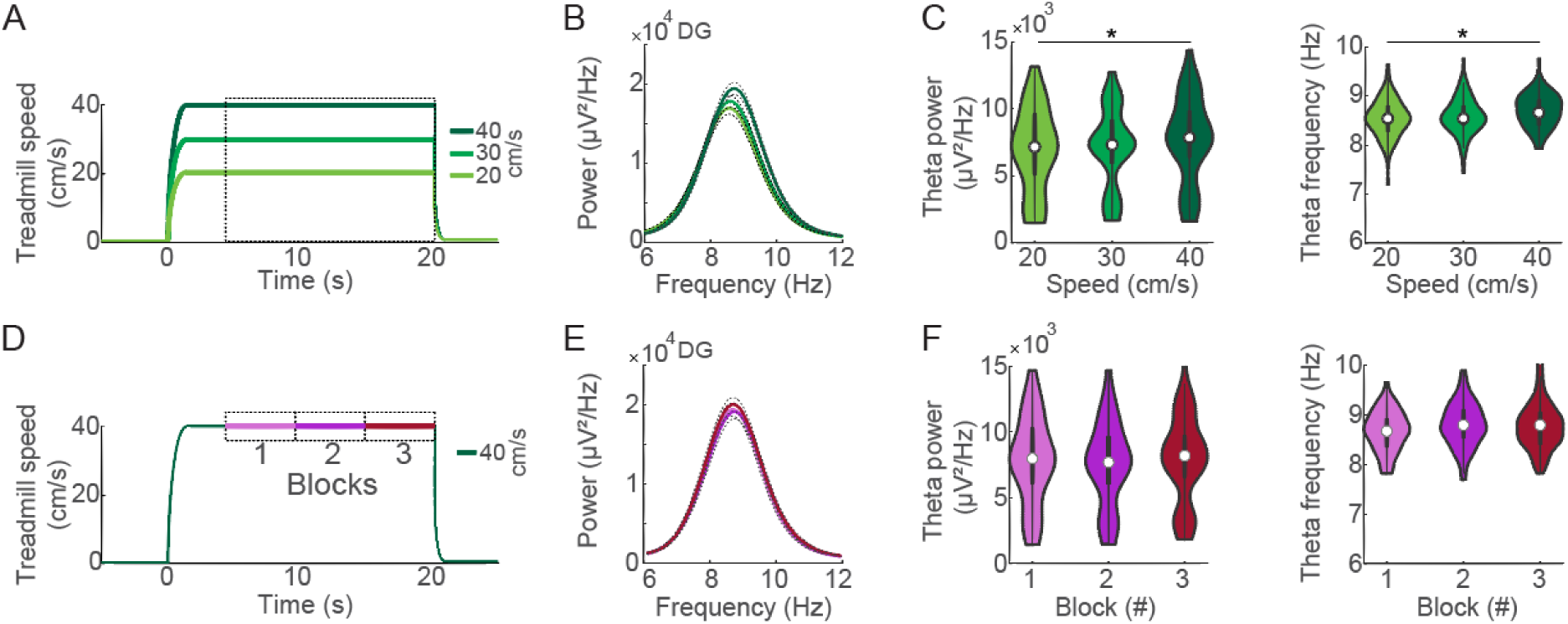
Theta power and frequency increase with treadmill running speed. (A)Treadmill protocols at constant running speeds of 20, 30, and 40 cm/s. To exclude movement transition effects, only running epochs between 5 and 20 s were analyzed. The dashed square depicts the interval of interest. (B) Mean power spectra at the theta band obtained from the left DG during treadmill running trials at 20, 30, and 40 cm/s. Solid lines represent means; dashed lines represent SEM. (C) Violin plots showing theta power (left) and theta peak frequency (right) during running at the three speeds. Asterisks denote p<0.05, Kruskal-Wallis test. (D) Treadmill protocol at a constant speed of 40 cm/s. For the subsequent analysis, the trials were divided into three consecutive 5-s blocks (5-10 s, 10-15 s, and 15-20 s). (E) Mean power spectra at the theta band from the left DG during each of the 5-s blocks. (F) Violin plots showing theta power (left) and theta peak frequency (right) during the 5-s blocks of runs at 40 cm/s.

### Treadmill acceleration rate does not modulate theta oscillations

We next compared theta oscillations during treadmill running at different acceleration rates (1, 1.5, and 2 cm/s^2^; Figure 5A). Spectral analysis revealed no significant differences in theta power or peak frequency in any of the hippocampal areas (e.g., as shown for the left DG in Figure 5B and C; see Supplementary Table 5-1). To control for the effect of instantaneous speed, we analyzed segments of the run where animals reached a similar speed (∼18–20 cm/s) under different acceleration rates (Figure 5D). Again, no significant differences in theta band power or peak frequency were observed (Figure 5E and F). Finally, we investigated whether progressive increases in speed within acceleration trials (2 cm/s^2^) modulated theta oscillations. To this end, we divided the last 15-s running period into three consecutive 5-s blocks (Figure 5G). This analysis revealed a significant increase in theta peak frequency across time blocks in DG, CA3, and CA1 areas of both hemispheres (e.g., shown for the left DG in Figure 5H and I; p < 0.01). In contrast, no significant changes in theta power were detected across the 5-s blocks in any hippocampal area (see Supplementary Table 5-1).

**Figure 5.**
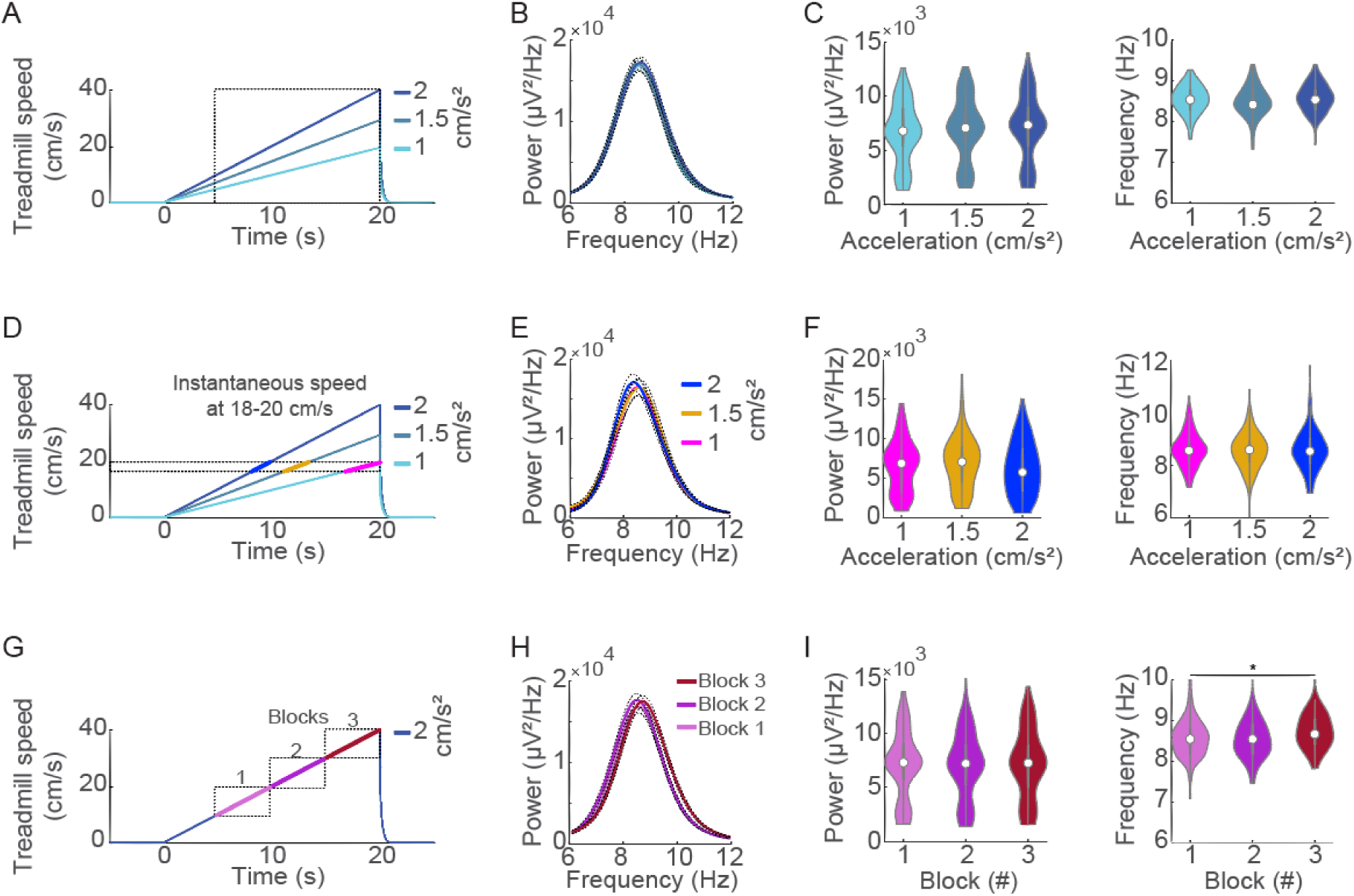
Theta power and frequency do not differ across accelerating running speeds. (A) Treadmill protocols at accelerating speeds of 1, 1.5, and 2 cm/s². Only running epochs between 5 and 20 s were analyzed. The dashed square depicts the interval of interest. (B) Mean power spectra at the theta band obtained from the left DG during runs at each acceleration level. (C) Violin plots showing theta power (left) and theta peak frequency (right) during accelerating running speeds. Asterisks denote p<0.05, Kruskal-Wallis test. (D) Same acceleration protocols as in A, but depicting intervals where the instantaneous speed was approximately 18–20 cm/s. (E) Mean power spectra at the theta band from the left DG during matched-speed intervals at different accelerations. (F) Violin plots of theta power (left) and peak frequency (right) at ∼18–20 cm/s across the three acceleration protocols. (G) Treadmill protocol at 2 cm/s^2^ acceleration, depicting the three 5-s blocks analyzed in H and I. (H) Mean power spectra at the theta band from the left DG across the three 5-s blocks. (I) Violin plots showing theta power (left) and peak frequency (right) across the three 5-s blocks. Asterisks indicate significant differences (p < 0.05, Kruskal–Wallis test).

## Discussion

We investigated how hippocampal theta oscillations relate to locomotion dynamics in rats using a controlled treadmill setup. Our findings reveal that movement initiation abruptly increases both theta power and frequency, but their subsequent dynamics diverge. Theta power remains elevated after movement onset and returns to baseline only after the treadmill stops. Theta frequency peaks shortly after movement begins and decays more rapidly, often returning toward pre-movement levels before the end of the running period (Figure 2). In addition, our results demonstrate that instantaneous running speed is the primary modulator of hippocampal theta activity. During steady treadmill running, both theta power and frequency were elevated compared to inter-trial rest periods (Figure 3). Increases in running speed led to direct increases in both theta power and frequency, with no additional changes observed during continuous running at a given constant speed (Figure 4). In contrast, the treadmill acceleration rate per se exhibited negligible influence on theta oscillations, with changes being primarily driven by the underlying instantaneous speed (Figure 5). The analysis of controlled speed conditions provides clear evidence on the distinct impacts of movement and reinforces the dominant role of instantaneous speed in governing hippocampal theta dynamics.

Previous reports have investigated the relationship between hippocampal oscillations and rat locomotor behavior during open-field exploration and various maze tasks (Bland and Vanderwolf, 1972; Whishaw and Vanderwolf, 1973; Vanderwolf and Heron, 1964; Rivas et al., 1996; Shin et al., 2001; Bouwman et al., 2005; Geisler et al., 2007; Hinman et al., 2011; Ahmed and Mehta, 2012; Long et al., 2014; Bender et al., 2015; de Lima and Belchior, 2023). These studies generally agree that hippocampal theta power and frequency correlate positively with locomotion speed. However, this relationship is often confounded with task-specific behavioral demands, such as motor parameters (e.g., movement initiation, acceleration), and cognitive or emotional factors (e.g., motivation, anxiety, decision making, learning, and memory processes) (Montgomery et al., 2009; Richard et al., 2013; Wells et al., 2013; Belchior et al., 2014; Young et al., 2021).

Instead of recording freely moving rats, we used a computer-controlled treadmill that allowed precise control over the timing and pace of locomotion. It is noteworthy that this approach limits the natural movement repertoire that a rat performs in naturalistic conditions, and also introduces biases, such as forced pacing or lack of self-motivated movement. Nevertheless, the treadmill provides a valuable framework for isolating specific locomotor parameters, such as the transitions from rest to movement and the precise control over constant or gradually increasing locomotion speeds. Moreover, this approach constraints spontaneous behaviors or cognitive loads, thereby ensuring a controlled experimental setting.

Our results generally confirm long-established observations by Vanderwolf and colleagues that theta oscillations increase at the onset of voluntary movements (Vanderwolf and Heron, 1964; Bland and Vanderwolf, 1972; Whishaw and Vanderwolf, 1973). We extend these findings with precise measures of movement intensity, anatomical specificity, and the independent evaluation of theta power and frequency. This approach revealed distinct dynamics associated with movement transitions versus sustained locomotion. Specifically, we found abrupt and transient increases in theta power and frequency when rats changed from rest to movement (Figure 2). Faster movement transitions led to larger increases in theta power and frequency. Importantly, these changes in both theta power and frequency during movement transitions were consistently observed across all hippocampal areas (DG, CA3, and CA1), suggesting a coordinated response to the initiation and cessation of locomotion within the hippocampal formation.

Kuo et al. (2011) showed that CA1 theta power and frequency exhibit distinct dynamics at the start and maintenance of treadmill runs at constant speed. They found that theta power increased and remained elevated throughout the 16-s run, while theta frequency peaked at movement onset, and gradually declined over the following seconds. Consistent with these observations, here we found that theta power remained elevated longer than theta frequency following movement initiation, and only returned to baseline levels after movement to rest transitions (Figure 2). However, direct comparisons revealed that both theta power and frequency are higher during the sustained period of treadmill running than during inter-trial rest intervals, especially at constant speed trials (Figure 3). Notably, theta power during inter-trial rest intervals may reflect arousal or anticipatory task engagement rather than a neutral baseline of hippocampal activity. Our results highlight different magnitudes of change in theta power and frequency, and further extend the findings of Kuo et al. (2011) to DG and CA3 areas of the rat hippocampus.

More recent studies examining the relationship between theta oscillations and locomotion reported divergent findings. First, Kropf et al. (2021) used a forced locomotion apparatus (a bottomless car along a linear track) to evaluate the effects of speed transitions, such as acceleration and deceleration, on the power and frequency of hippocampal theta oscillations. The authors concluded that the instantaneous running speed per se did not modulate theta oscillations. Instead, they suggested that acceleration has a major influence on the modulation of theta frequency, with minor effects on theta power. A subsequent study by Kennedy et al. (2022) found that acceleration exerted only a minor influence on theta frequency and power, whereas the instantaneous running speed accounted for most changes in theta frequency in freely moving rats performing a maze task. Furthermore, in an attempt to disentangle the incompatible results, Kennedy et al. (2022) also reevaluated the dataset collected by Kropff et al. (2021) and argued that the original conclusions were likely affected by biased sampling of speed and acceleration within the speed-clamping task. Kennedy et al. (2022) reanalyzed the data, corrected the sampling distribution, and confirmed that speed remained the dominant factor even during acceleration and deceleration.

Our findings support Kennedy et al. (2022) by showing that theta frequency was primarily determined by the instantaneous speed of locomotion. Theta frequency changed with speed in all hippocampal areas and at both constant and acceleration trials, i.e., across different speeds in steady running conditions (Figure 4) and accompanying running speeds at acceleration trials (Figure 5). In contrast, theta power was only modestly affected by speed in conditions of steady treadmill running. This dissociation is notable: while instantaneous speed robustly drives theta frequency increases across the entire DG-CA3-CA1 axis, the modulation of theta power by speed appears to be more prominent in the input (DG) and associative (CA3) areas of the hippocampus compared to the primary output area (CA1). Although this regional difference may reflect distinct computations or levels of network engagement, it is also possible that the observed effects may arise from statistical variability rather than true lateralized or subfield-specific mechanisms. In addition, we show that neither theta power nor frequency changed across the time of running at a given constant speed (Figure 4). These results suggest that running speed has a major influence on theta frequency and a limited effect on theta power.

In contrast to the findings of Kropff et al. (2021), our results indicate that acceleration does not significantly influence hippocampal theta oscillations. For instance, we observed that different acceleration rates on the treadmill did not modulate theta power or frequency (Figure 5), even under tightly controlled conditions. For instance, time intervals characterized by similar instantaneous running speeds but distinct acceleration rates presented no differences in theta power and frequency. Although our study focused on a specific range of acceleration rates relevant to controlled treadmill locomotion, these findings provide compelling evidence that acceleration is not a major driver of theta oscillations and reinforce instantaneous speed as the dominant modulating factor.

It is important to acknowledge the limitations of our study. First, recordings were restricted to the dorsal hippocampus (DG, CA3, and CA1), leaving open the possibility that theta modulation differs along the septotemporal axis or in extrahippocampal regions. Second, the relatively narrow range of locomotion speeds tested does not rule out the emergence of nonlinear effects at more extreme velocities, which could engage additional mechanisms such as fatigue or exhaustion of the animals. Finally, as our analyses focused on LFPs, the cellular and circuit-level mechanisms linking speed, movement transitions, and theta generation remain unresolved. Future studies incorporating simultaneous single-unit recordings or optogenetic manipulations will be necessary to uncover causal relationships within the underlying hippocampal circuitry.

In summary, movement transitions exert a strong influence on hippocampal theta oscillations, with distinct temporal dynamics for power and frequency. Steady periods of treadmill running yield elevated theta power and frequency relative to rest. Across running speeds, theta frequency—and to a lesser extent, theta power—tracks instantaneous speed. Our findings demonstrate that the combination of movement transitions and instantaneous locomotion speed, rather than acceleration rate per se, governs theta activity in the hippocampus. This speed-dependent modulation of theta may, in turn, shape other hippocampal computations, including the activity of place, time, and speed cells (Geisler et al., 2007; Eichenbaum, 2014; Góis and Tort, 2018).

## Acknowledgements

This research was supported in part by the Coordenação de Aperfeiçoamento de Pessoal de Nível Superior (CAPES), and by the Conselho Nacional de Desenvolvimento Científico e Tecnológico (CNPq).

