## Supplementary Materials for "HIPPOCAMPAL THETA OSCILLATIONS DURING CONTROLLED SPEED RUNS ON A TREADMILL"

| P-values for theta power increases at locomotion onset | | | | | | |
|---|---|---|---|---|---|---|
| Condition / Area | Left DG | Right DG | Left CA3 | Right CA3 | Left CA1 | Right CA1 |
| S1 = 20 cm/s | $1.0x10^{-15}$* | $1.0x10^{-15}$* | $1.0x10^{-15}$* | $1.0x10^{-15}$* | $2.0x10^{-14}$* | $1.3x10^{-11}$* |
| S2 = 30 cm/s | $1.0x10^{-15}$* | $1.0x10^{-15}$* | $1.0x10^{-15}$* | $1.0x10^{-15}$* | $1.0x10^{-15}$* | $4.8x10^{-13}$* |
| S3 = 40 cm/s | $1.0x10^{-15}$* | $1.0x10^{-15}$* | $1.0x10^{-15}$* | $1.0x10^{-15}$* | $1.0x10^{-14}$* | $3.5x10^{-13}$* |
| A1 = 1.0 cm/s² | $1.0x10^{-10}$* | $1.0x10^{-10}$* | $3.6x10^{-8}$* | $4.3x10^{-6}$* | $1.0x10^{-10}$* | $1.0x10^{-10}$* |
| A2 = 1.5 cm/s² | $1.0x10^{-10}$* | $1.0x10^{-10}$* | $1.0x10^{-10}$* | $1.0x10^{-10}$* | $1.0x10^{-10}$* | $1.0x10^{-10}$* |
| A3 = 2.0 cm/s² | $1.0x10^{-10}$* | $1.0x10^{-10}$* | $1.0x10^{-10}$* | $1.0x10^{-10}$* | $1.0x10^{-10}$* | $1.0x10^{-10}$* |

| P-values for theta frequency increases at locomotion onset | | | | | | |
|---|---|---|---|---|---|---|
| Condition / Area | Left DG | Right DG | Left CA3 | Right CA3 | Left CA1 | Right CA1 |
| S1 = 20 cm/s | $1.0x10^{-6}$* | $1.0x10^{-8}$* | $1.0x10^{-6}$* | $1.1x10^{-4}$* | $1.0x10^{-8}$* | $1.0x10^{-8}$* |
| S2 = 30 cm/s | $1.0x10^{-8}$* | $1.0x10^{-8}$* | $1.0x10^{-8}$* | $8.7x10^{-5}$* | $1.0x10^{-8}$* | $1.0x10^{-8}$* |
| S3 = 40 cm/s | $1.0x10^{-8}$* | $1.0x10^{-8}$* | $1.0x10^{-8}$* | $1.1x10^{-5}$* | $1.0x10^{-6}$* | $3.0x10^{-6}$* |
| A1 = 1.0cm/s² | $1.0x10^{-5}$* | $1.0x10^{-5}$* | $1.0x10^{-5}$* | $1.0x10^{-5}$* | $1.0x10^{-5}$* | $1.0x10^{-5}$* |
| A2 = 1.5 cm/s² | $1.0x10^{-5}$* | $3.0x10^{-4}$* | $8.0x10^{-4}$* | $3.3x10^{-3}$* | $5.0x10^{-4}$* | $1.1x10^{-3}$* |
| A3 = 2.0 cm/s² | $6.0x10^{-4}$* | $1.0x10^{-5}$* | $1.0x10^{-4}$* | $1.0x10^{-4}$* | $1.0x10^{-5}$* | $1.0x10^{-5}$* |

**Supplementary Table 2-1.** P-values for theta power (top table) and theta frequency (bottom table) comparing inter-trial rest intervals to movement onset across constant- and accelerating-speed protocols. Each value reflects the result of a Wilcoxon signed-rank sum test applied to 2-second epochs before and after treadmill onset. Asterisks indicate statistically significant comparisons ($p < 0.05$).

| P-values for theta power decreases at locomotion offset | | | | | | |
|---|---|---|---|---|---|---|
| Condition / Area | Left DG | Right DG | Left CA3 | Right CA3 | Left CA1 | Right CA1 |
| S1 = 20 cm/s | $1.0 \times 10^{-8}$* | $1.0 \times 10^{-8}$* | $1.0 \times 10^{-8}$* | $1.0 \times 10^{-8}$* | $1.0 \times 10^{-7}$* | $7.9 \times 10^{-5}$* |
| S2 = 30 cm/s | $1.0 \times 10^{-8}$* | $1.0 \times 10^{-8}$* | $1.0 \times 10^{-8}$* | $1.0 \times 10^{-8}$* | $1.0 \times 10^{-8}$* | $3.9 \times 10^{-4}$* |
| S3 = 40 cm/s | $1.0 \times 10^{-8}$* | $1.0 \times 10^{-8}$* | $1.0 \times 10^{-8}$* | $1.0 \times 10^{-8}$* | $1.0 \times 10^{-8}$* | $3.0 \times 10^{-7}$* |
| A1 = 1.0 cm/s² | $1.0 \times 10^{-8}$* | $1.0 \times 10^{-8}$* | $1.0 \times 10^{-8}$* | $1.0 \times 10^{-8}$* | $1.0 \times 10^{-8}$* | $4.0 \times 10^{-4}$* |
| A2 = 1.5 cm/s² | $1.0 \times 10^{-8}$* | $1.0 \times 10^{-8}$* | $1.0 \times 10^{-8}$* | $1.0 \times 10^{-8}$* | $1.0 \times 10^{-8}$* | $1.2 \times 10^{-5}$* |
| A3 = 2.0 cm/s² | $1.0 \times 10^{-8}$* | $1.0 \times 10^{-8}$* | $1.0 \times 10^{-8}$* | $1.0 \times 10^{-8}$* | $1.0 \times 10^{-8}$* | $2.2 \times 10^{-6}$* |

| P-values for theta frequency decreases at locomotion offset | | | | | | |
|---|---|---|---|---|---|---|
| Condition / Area | Left DG | Right DG | Left CA3 | Right CA3 | Left CA1 | Right CA1 |
| S1 = 20 cm/s | $1.0 \times 10^{-5}$* | $1.0 \times 10^{-5}$* | $1.0 \times 10^{-5}$* | $1.0 \times 10^{-5}$* | $1.0 \times 10^{-5}$* | $1.0 \times 10^{-5}$* |
| S2 = 30 cm/s | $1.0 \times 10^{-2}$* | $1.5 \times 10^{-3}$* | $8.5 \times 10^{-3}$* | $1.0 \times 10^{-5}$* | $1.0 \times 10^{-4}$* | $3.0 \times 10^{-4}$* |
| S3 = 40 cm/s | $1.0 \times 10^{-4}$* | $1.1 \times 10^{-1}$ | $1.0 \times 10^{-5}$* | $1.0 \times 10^{-5}$* | $1.0 \times 10^{-4}$* | $1.0 \times 10^{-4}$* |
| A1 = 1.0 cm/s² | $1.0 \times 10^{-4}$* | $1.0 \times 10^{-5}$* | $1.0 \times 10^{-5}$* | $3.0 \times 10^{-4}$* | $1.0 \times 10^{-4}$* | $5.0 \times 10^{-4}$* |
| A2 = 1.5 cm/s² | $1.0 \times 10^{-3}$* | $1.3 \times 10^{-2}$* | $1.0 \times 10^{-5}$* | $1.0 \times 10^{-4}$* | $4.7 \times 10^{-2}$* | $1.4 \times 10^{-1}$ |
| A3 = 2.0 cm/s² | $1.0 \times 10^{-4}$* | $1.0 \times 10^{-5}$* | $1.0 \times 10^{-5}$* | $1.0 \times 10^{-5}$* | $1.0 \times 10^{-5}$* | $1.0 \times 10^{-5}$* |

**Supplementary Table 2-2.** P-values for theta power (top table) and theta frequency (bottom table) comparing movement offset to inter-trial rest intervals across constant- and accelerating-speed protocols. Each value reflects the result of a Wilcoxon signed-rank sum test applied to 2-second epochs before and after treadmill stop. Asterisks indicate statistically significant comparisons ($p < 0.05$).

| P-values for theta power: sustained treadmill running vs. inter-trial rest | | | | | | |
|---|---|---|---|---|---|---|
| Condition / Area | Left DG | Right DG | Left CA3 | Right CA3 | Left CA1 | Right CA1 |
| S1 = 20 cm/s | $1.0 \times 10^{-5}$* | $1.0 \times 10^{-5}$* | $1.0 \times 10^{-5}$* | $1.0 \times 10^{-5}$* | $1.0 \times 10^{-5}$* | $2.0 \times 10^{-4}$* |
| S2 = 30 cm/s | $1.0 \times 10^{-5}$* | $1.0 \times 10^{-5}$* | $1.0 \times 10^{-5}$* | $1.0 \times 10^{-5}$* | $1.0 \times 10^{-5}$* | $2.1 \times 10^{-3}$* |
| S3 = 40 cm/s | $1.0 \times 10^{-5}$* | $1.0 \times 10^{-5}$* | $1.0 \times 10^{-5}$* | $1.0 \times 10^{-5}$* | $1.0 \times 10^{-5}$* | $1.0 \times 10^{-5}$* |
| A1 = 1.0 cm/s² | $1.0 \times 10^{-5}$* | $1.0 \times 10^{-5}$* | $1.0 \times 10^{-5}$* | $1.0 \times 10^{-5}$* | $1.0 \times 10^{-5}$* | $1.7 \times 10^{-3}$* |
| A2 = 1.5 cm/s² | $1.0 \times 10^{-5}$* | $1.0 \times 10^{-5}$* | $1.0 \times 10^{-5}$* | $1.0 \times 10^{-5}$* | $1.0 \times 10^{-5}$* | $9.0 \times 10^{-4}$* |
| A3 = 2.0 cm/s² | $1.0 \times 10^{-5}$* | $1.0 \times 10^{-5}$* | $1.0 \times 10^{-5}$* | $1.0 \times 10^{-5}$* | $1.0 \times 10^{-5}$* | $6.0 \times 10^{-4}$* |

| P-values for theta frequency: sustained treadmill running vs. inter-trial rest | | | | | | |
|---|---|---|---|---|---|---|
| Condition / Area | Left DG | Right DG | Left CA3 | Right CA3 | Left CA1 | Right CA1 |
| S1 = 20 cm/s | $3.6 \times 10^{-3}$* | $1.6 \times 10^{-3}$* | $7.5 \times 10^{-3}$* | $4.0 \times 10^{-3}$* | $2.0 \times 10^{-4}$* | $1.0 \times 10^{-4}$* |
| S2 = 30 cm/s | $7.7 \times 10^{-3}$* | $1.8 \times 10^{-2}$* | $5.6 \times 10^{-3}$* | $4.0 \times 10^{-2}$* | $3.7 \times 10^{-3}$* | $9.0 \times 10^{-4}$* |
| S3 = 40 cm/s | $3.9 \times 10^{-2}$* | $3.3 \times 10^{-2}$* | $5.8 \times 10^{-1}$ | $3.2 \times 10^{-1}$ | $1.1 \times 10^{-2}$* | $2.0 \times 10^{-2}$* |
| A1 = 1.0 cm/s² | $1.0 \times 10^{-5}$* | $1.0 \times 10^{-5}$* | $4.0 \times 10^{-4}$* | $7.0 \times 10^{-4}$* | $1.0 \times 10^{-5}$* | $1.0 \times 10^{-5}$* |
| A2 = 1.5 cm/s² | $4.2 \times 10^{-1}$ | $3.4 \times 10^{-1}$ | $1.5 \times 10^{-1}$ | $5.4 \times 10^{-2}$ | $5.6 \times 10^{-1}$ | $4.9 \times 10^{-1}$ |
| A3 = 2.0 cm/s² | $1.9 \times 10^{-1}$ | $3.0 \times 10^{-1}$ | $6.2 \times 10^{-1}$ | $9.9 \times 10^{-1}$ | $1.0 \times 10^{-1}$ | $1.8 \times 10^{-1}$ |

**Supplementary Table 3-1.** P-values for theta power (top table) and theta frequency (bottom table) comparing treadmill running trials to inter-trial rest intervals across constant- and accelerating-speed protocols. Each comparison was based on 15-second running epochs (excluding the first 5 seconds following movement onset) and matched 5-second rest intervals. Asterisks indicate statistically significant differences ($p < 0.05$) based on the Wilcoxon signed-rank sum test.

| Statistical comparison of theta power and frequency across constant-speed protocols | | | | | | |
|---|---|---|---|---|---|---|
| Condition / Area | Left DG | Right DG | Left CA3 | Right CA3 | Left CA1 | Right CA1 |
| Theta power | $3.5x10^{-2}$* | $3.7x10^{-1}$ | $3.1x10^{-1}$ | $5.0x10^{-3}$* | $9.1x10^{-1}$ | $2.7x10^{-1}$ |
| Theta frequency | $2.0x10^{-3}$* | $1.0x10^{-5}$* | $1.3x10^{-2}$* | $1.0x10^{-5}$* | $3.0x10^{-3}$* | $2.0x10^{-3}$* |

| Statistical comparison of theta power and frequency across 5-s blocks during steady-state running at 40 cm/s | | | | | | |
|---|---|---|---|---|---|---|
| Condition / Area | Left DG | Right DG | Left CA3 | Right CA3 | Left CA1 | Right CA1 |
| Theta power | $9.1x10^{-1}$ | $9.3x10^{-1}$ | $7.8x10^{-1}$ | $8.6x10^{-1}$ | $9.3x10^{-1}$ | $9.7x10^{-1}$ |
| Theta frequency | $1.6x10^{-1}$ | $3.6x10^{-1}$ | $8.2x10^{-1}$ | $9.9x10^{-1}$ | $3.8x10^{-1}$ | $3.8x10^{-1}$ |

**Supplementary Table 4-1.** P-values for theta power and theta peak frequency differences across treadmill runs at constant speeds of 20, 30, and 40 cm/s (top table), and across consecutive 5-s blocks (5-10 s, 10-15 s, and 15-20 s) during runs at 40 cm/s (bottom table). Each value reflects the result of a Kruskal–Wallis test across conditions. Asterisk denotes statistical significance ($p<0.05$).

| Statistical comparison of theta power and frequency across acceleration protocols | | | | | | |
|---|---|---|---|---|---|---|
| Condition / Area | Left DG | Right DG | Left CA3 | Right CA3 | Left CA1 | Right CA1 |
| Theta power | $1.8 \times 10^{-1}$ | $3.5 \times 10^{-1}$ | $3.1 \times 10^{-1}$ | $2.5 \times 10^{-1}$ | $7.5 \times 10^{-1}$ | $8.5 \times 10^{-1}$ |
| Theta frequency | $1.8 \times 10^{-1}$ | $2.2 \times 10^{-1}$ | $5.9 \times 10^{-1}$ | $4.2 \times 10^{-1}$ | $3.6 \times 10^{-1}$ | $2.4 \times 10^{-1}$ |
| **Theta power and frequency comparisons at equal speeds but different accelerations** | | | | | | |
| Condition / Area | Left DG | Right DG | Left CA3 | Right CA3 | Left CA1 | Right CA1 |
| Theta power | $3.4 \times 10^{-1}$ | $2.3 \times 10^{-1}$ | $4.4 \times 10^{-1}$ | $8.3 \times 10^{-1}$ | $7.9 \times 10^{-1}$ | $9.7 \times 10^{-1}$ |
| Theta frequency | $8.4 \times 10^{-1}$ | $9.5 \times 10^{-1}$ | $6.4 \times 10^{-1}$ | $8.2 \times 10^{-1}$ | $9.5 \times 10^{-1}$ | $8.5 \times 10^{-1}$ |
| **Statistical comparison of 5-s blocks during acceleration trials at 2 cm/s²** | | | | | | |
| Condition / Area | Left DG | Right DG | Left CA3 | Right CA3 | Left CA1 | Right CA1 |
| Theta power | $8.2 \times 10^{-1}$ | $8.8 \times 10^{-1}$ | $4.8 \times 10^{-1}$ | $4.6 \times 10^{-1}$ | $5.6 \times 10^{-1}$ | $7.0 \times 10^{-1}$ |
| Theta frequency | $1.0 \times 10^{-3}$* | $1.0 \times 10^{-3}$* | $4.0 \times 10^{-3}$* | $5.0 \times 10^{-3}$* | $8.0 \times 10^{-3}$* | $2.8 \times 10^{-3}$* |

**Supplementary Table 5-1.** P-values for theta power and theta frequency during accelerating running speeds. The upper table shows comparisons across acceleration trials at 1, 1.5, and 2 cm/s². The central table shows comparisons across treadmill running trials at the same instantaneous speed (18-20 cm/s) but at distinct acceleration rates. The lower table shows comparisons across 5-s blocks of treadmill running trials during accelerating speeds at 2 cm/s². Asterisks indicate statistically significant differences ($p < 0.05$) based on the Kruskal–Wallis test.
